# Phosphorylation of JIP4 at S730 presents anti-viral properties against influenza A virus infection

**DOI:** 10.1101/2021.01.22.427772

**Authors:** Juliana Del Sarto, Vanessa Gerlt, Marcel Edgar Friedrich, Darisuren Anhlan, Viktor Wixler, Mauro Martins Teixeira, Yvonne Boergeling, Stephan Ludwig

**Author notes:** shared senior authorship (Yvonne Boergeling, Stephan Ludwig).

## Abstract

Influenza A virus (IAV) is the causative agent of flu disease that results in annual epidemics and occasional pandemics. IAV alters several signaling pathways of the cellular host response in order to promote its replication. Therefore, our group investigates different host cell pathways modified in IAV infection as promising targets for long-lasting therapeutic approaches. Here, we show that c-Jun NH2-terminal kinase (JNK)-interacting protein (JIP) 4 is dynamically phosphorylated in IAV infection. Lack of JIP4 resulted in higher virus titers with significant differences in viral protein and mRNA accumulation as early as within the first replication cycle. In accordance, decreased IAV titers and protein accumulation was observed during overexpression of JIP4. Strikingly, the anti-viral function of JIP4 does neither originate from a modulation of JNK or p38 MAPK pathways, nor from altered expression of interferons or interferon-stimulated genes, but rather from a direct reduction of viral polymerase activity. Furthermore, interference of JIP4 with IAV replication is linked to phosphorylation of the serine at position 730, that is sufficient to impede with the viral polymerase and is mediated by the Raf/MEK/ERK pathway. Collectively, we provide evidence that JIP4, a host protein modulated in IAV infection, exhibits anti-viral properties that are dynamically controlled by its phosphorylation at S730.

## Introduction

Influenza A virus (IAV) is a common respiratory pathogen that infects up to 10% of the global population annually. IAV shows high mutation rates that lead to a constant *antigenic drift*. Furthermore, there is a permanent threat of reassortment, potentially leading to the generation of new highly pathogenic strains (*antigenic shift*). As a result, vaccines have to be annually updated according to predictions of circulating strains for the next epidemic season. Additionally, antiviral drugs that target IAV proteins present limited long-term efficiency, due to increasing viral resistance. Most of the FDA-approved IAV antivirals used in the clinics today, act as neuraminidase inhibitors (Zanamivir, Oseltamivir and Peramivir) or M2 channel blockers (Adamantane), all targeting specific viral proteins (1). Unfortunately, high rates of resistance against these drugs have been documented. Important examples are the global spread of oseltamivir-resistant seasonal A (H1N1) viruses in 2007, and adamantane-resistant pandemic A (H1N1) viruses in 2009 (2). As a strategy to avoid viral resistance, we aim to interfere with host cell proteins and pathways, posing less pressure for the development of viral escape variants.

During viral infection, activity of several host cell signaling pathways is upregulated. While this can be primarily seen as a defense response of the infected cell, viruses have acquired the capability to misuse some of these pathways for their own purpose. Different pathways have already been shown to act virus-supportive such as the Raf/MEK/ERK cascade that modulates the export of viral ribonucleoproteins (RNPs) from the nucleus into the cytoplasm in later stages of the IAV replication cycle (3). Interestingly, another member of the family of mitogen-activated protein kinases (MAPK), the c-Jun NH2-terminal kinase (JNK), also showed virus-supportive activities (4) although it has been implicated in the regulation of transcription factor activator protein 1 (AP-1) that is needed for efficient host antiviral responses (5). Likewise, activation of MAPK p38 could directly be linked to the expression of interferon (IFN) and pro-inflammatory cytokines as well as to the modulation of IFN-induced responses (6). Different proteins have been reported to regulate or to be regulated by the pathways mentioned above, including a newly discovered protein named JNK-interacting protein 4 (JIP4) (7).

Besides three similar proteins JIP1, JIP2 and JIP3, JIP4 belongs to the group of JNK-interacting proteins. All proteins of this group are known scaffold proteins that can interact with JNK and kinesin light chain (7). JIP4, also known as sperm associated antigen 9 (SPAG9) or JNK– associated leucine zipper protein (JLP), is the most recently discovered protein of the JIP group, resulting in still little knowledge about its functions. Limited amounts of studies have been published identifying different activities of JIP4 in different scenarios. One showed that JIP4 phosphorylation is important during G2 phase of mitosis by its interaction with p38 MAPK (8). JIP4 has also been correlated with the promotion of an invasive phenotype of breast cancer cells through its interaction with proteins important for endosome tubulation and exocytosis (9). Despite the close relation of JIP4 to JNK and p38 MAPKs, no functional association between JIP4 and virus infection has been described so far.

Therefore, we aimed to understand the role of JIP4 in IAV infection. In this study, we report anti-viral properties of JIP4 against IAV *in vitro*. JIP4 phosphorylation at S730 interferes with viral polymerase activity resulting in decreased viral replication. Hence, JIP4 represents a potential host target for therapeutic approaches.

## Methods

### Virus strains and cell lines

IAV H1N1 strains A/PuertoRico/8/34, A/WSN/33 and A/Hamburg/04/09 were taken from virus collections of the Institute of Virology Muenster. A/Thailand/KAN-1/2004 (H5N1) was used with kind permission from P. Puthavathana (Bangkok, Thailand). A/FPV/79/Bratislava (H7N7, fowl plague virus) was originally obtained from the Institute of Virology in Giessen, Germany. Influenza viruses as well as vesicular stomatitis virus strain Indiana (VSV) were propagated on Madin-Darby canine kidney (MDCKII) cells cultured in minimal essential medium (MEM-Sigma) containing 10% (v/v) FCS (Biochrom). Human alveolar epithelial cells (A549) and green monkey epithelial cells (Vero) were cultured in DMEM (Sigma) containing 10% FCS. All cell lines were originally purchased from ATCC and have been passaged in the laboratory. At regular intervals cells are checked for their identity by SNP-profiling (Multiplexion).

### Reagents and plasmids

JIP4-expressing plasmids pRP-EGFP/Puro-CAG-JIP4 wild type (WT), JIP4 S730A and JIP4 S730E were purchased from VectorBuilder. All constructs for the viral polymerase reconstitution assay were created as previously described (10). All plasmids were transformed in competent bacterial cells (XL-Gold), propagated in LB medium with 25 μg/ml ampicillin and plasmid purifications were performed according to the manufacturers’ instructions (Sigma).

Recombinant human type I interferon β (IFNβ) was obtained from pbl assay science and added in a concentration of 100 units/ml to the medium of seeded A549 cells. For the inhibition of the Raf/MEK/ERK pathway, Vero cells were incubated with 10 μM of the MEK-specific inhibitor U0126 (DMSO soluble, Promega).

### Mass spectrometry

For the measurement of JIP4 phosphorylation dynamics, stable isotope labelling by amino acids in cell culture (SILAC) was used. Human A549 cells stably labelled with either ‘light’ lysine (12C6, 14N2) and arginine (12C6, 14N4), ‘medium’ lysine (13C6, 14N2) and arginine (13C6, 14N2) or ‘heavy’ lysine (13C6, 15N2) and arginine (13C6, 15N4) were infected with PR8, FPV or KAN-1 (MOI = 5) for 0, 2, 4, 6 and 8 h. Here, ‘medium’‐labelled and ‘heavy’‐labelled cells were infected for 2 or 6 h and 4 or 8 h, respectively, while ‘light’‐labelled cells were always used as non‐infected controls (0 h). Lysates from cells infected for 0, 2 and 4 h were mixed equally and used as first sample, while equally mixed lysates from cells infected for 0, 6 and 8 h were used as second sample. After tryptic digestion, samples were subjected to cation exchange chromatography and TiO2‐based phosphopeptide enrichment chromatography followed by LC‐MS/MS analysis on a Proxeon Easy‐nLC coupled to an LTQ‐Orbitrap XL mass spectrometer. Data were processed using Mascot and MaxQuant software (v1.2.2.9) as previously described (11, 12). Dynamics in phosphorylation intensities of JIP4-derived peptides were quantified in relation to the ratio of total JIP4 protein expressed at the different time points analyzed, normalized to intensities in non-infected cells.

### Transfection and RNA stimulation

For JIP4 silencing, A549 cells were transfected in suspension with 50 pmol per 3×10^5^ cells of scrambled siRNAs (control, 5’UUCUCCGAACGUGUCACGU3’) or siRNAs specific for JIP4 (5’GAGCAUGUCUUUACAGAUCUU3’) using the transfection reagent Lipofectamine^R^ 2000 (Invitrogen) according to manufacturers’ instructions. Medium was exchanged 24 hours post transfection (hpt) and infection was carried out for 48 h.

For overexpression of WT JIP4 or JIP4 mutants, A549 or Vero cells were transfected with 3 μg of plasmids with transfection reagent Lipofectamine^R^ 2000 (Invitrogen) according to manufacturers’ instructions 24 h post seeding. Empty vector plasmids were used as transfection control. Medium exchange was performed 4 hpt, infection was carried out for 24 h.

For the production of control RNA (cRNA) and viral RNA (vRNA), A549 cells were either infected with 5 MOI of PR8 for 8 h (vRNA) or mock-infected with PBS (cRNA) and cells were lysed for RNA isolation. RNAs were then stored at −80°C until usage. For stimulation with RNA, cells were transfected with c/vRNA as described previously for siRNA transfections. Cells were lysed 3 hpt and proceeded for Western Blot analyses.

### Virus titration by standard plaque assay

Virus-containing supernatants were collected at specified time points and stored at −80°C. Viral titers were assessed by standard plaque assays as described previously (13). Briefly, monolayers of MDCKII cells were infected with serial dilutions of IAV-containing supernatants for 30 min at 37°C to allow for virus adsorption. The viral inoculum was subsequently aspirated and replaced by 2 ml of plaque medium (0.2% bovine serum albumin, 1 mM MgCl_2_, 0.9 mM CaCl_2_, 100 U/ml penicillin, 0.1 mg/ml streptomycin, 0.3% DEAE-Dextran and 1.5% NaHCO_3_) supplemented with 0.6% Agar (Oxoid). Plates were maintained at 37°C with 5% CO_2_ for 3 days until manual counting of plaques. All results are expressed as plaque forming units (pfu)/ml.

### RNA extraction, cDNA synthesis and quantitative real-time PCR

A549 cells were infected with PR8 at a MOI of 5 and lysates were collected every two hours post infection (hpi) until 8 h RNA extraction was performed with Monarch total RNA Miniprep kit (New England Biolabs GmbH) according to manufacturers’ instructions. cDNA synthesis was performed with 1 μg of RNA and 0.5 μg of oligo-d(T) primers, which was incubated for 5 min at 70°C, followed by 1 min at 4°C and 2 min at 37°C. The MasterMix (4 μl of 5x buffer; 2 μl [10 mM] dNTP; 0.5 μl Revert Aid H minus reverse transcriptase) were added per sample as instructed by Thermofisher Scientific. The samples were then incubated for 1 h at 42°C followed by 10 min at 70°C and finally stored at −20°C until qRT-PCR assay. qRT-PCR was performed with 4 μl Brilliant III Ultra-Fast SYBR QPCR as instructed by the manufacturer (Agilent Santa Clara). All primers for quantification of either expression of viral m/cRNAs or antiviral responses are shown below.

Primers for analysis of viral mRNAs: M1 sense 5’tgcaaaaacatcttcaagtctctg 3’ and anti-sense 5’agatgagtcttctaaccgaggtcg 3’; PB1 sense 5’catacagaagaccagtcgggat 3’ and anti-sense 5’gtctgagctcttcaatggtgga 3’; NS sense 5’gaggacttgaatggaatgataaca 3’ and anti-sense 5’gtctcaattcttcaatcaatcaaccatc 3’; housekeeping gene GAPDH sense 5’gcaaattccatggcaccgt 3’ and anti-sense 5’gccccacttgatttggagg 3’; and SPAG9 sense 5’gcttttgatcgcaatacagaatctc 3’and anti-sense 5’aacttcccgacccattcctagt 3’.

Primers used for analyses of antiviral responses: IFNβ sense 5’gcgctcagtttcggaggtaacctgt 3’ and anti-sense 5’ggccatgaccaagtgtctcctcc 3’; MxA sense 5’gaagggcaactcctgacagt 3’ and anti-sense 5’ gtttccgaagtggacatcgca 3’; IP10 sense 3’ggaacctccagtctcagcacca 3’ and anti-sense 5’agacatctcttctcacccttc 3’; and OAS1 sense 5’ ctggccaggtggaaggag 3’ and anti-sense 5’agctacctcggaagcacctt 3’.

### Strand-specific quantitative real-time PCR

The protocol and primers for cDNA synthesis and qRT-PCR were used as described previously (14). Briefly, 750 ng of RNA diluted in a total volume of 3.5 μl was incubated with 0.5 μl Oligo-d(T) and viral strand-specific primers for vRNA, cRNA and mRNA. Samples were then incubated for 10 min at 65°C, 5 min at 4°C and 5 min at 60°C before addition of preheated MasterMix (4 μl First strand Buffer [5x], 1 μ1 DTT, 1 μl [10mM] dNTP, 0.5 μl Revert Aid Premium RT (Maxima) and 8 μl ddH2O) according to the manufacturer (Thermofisher). Next, samples were incubated for 1 h at 60°C followed by 5 min at 85°C before storage at −20°C until qRT-PCR analyses. qRT-PCR was performed as explained in the section above.

### Western Blot analysis

Cells were collected with radioimmunoprecipitation assay (RIPA) lysis buffer supplemented with protease and phosphatase inhibitors every two hours after infection until 8 hpi and kept at −20°C. Samples were centrifuged and relative protein amounts were determined by Bradford assay (Protein Assay dye Reagent Concentrate – BIORAD). Subsequently, samples were diluted 5:1 in Laemmli buffer, separated by SDS-PAGE and blotted using nitrocellulose membranes. Samples were analyzed for the expression of the viral proteins polymerase basic protein 1 (PB1), non-structural protein 1 (NS1) and matrix protein 1 (M1) (Genetex) or cellular proteins such as tubulin, JIP4, p38 or p-p38 MAPK T180/Y182 (Cell Signaling Technology), pJNK pT183/pY185 (BD Bioscience), JNK (Cell Signaling Technology).

### Viral polymerase activity

Viral polymerase activity was measured in a mini-genome assay as described previously (10). Briefly, Vero cells were transfected with a plasmid mixture containing influenza A virus polymerase subunits PB1, polymerase basic protein 2 (PB2), polymerase acidic protein (PA) (200 ng of each) and nucleoprotein (NP) (400 ng), as well as with a polymerase I-driven expression plasmid (25 ng) expressing an influenza virus-like RNA coding for the reporter protein firefly luciferase (vRNA-luciferase) and plasmid coding for renilla luciferase (25 ng) used for normalization to transfection efficiencies. JIP4 (3 μg) was overexpressed together with the previously described plasmids. The empty vector (EV, 3 μg) was used as positive control. As a negative control, PB1 plasmid was excluded from the mix. Medium was exchanged for DMEM 10% FCS 4 hpt and cells were collected after 24 h for measurement of luciferase activity by Dual Luciferase assay (promega). Relative light units (RLU) were normalized to respective Renilla activity and are expressed in percentage over positive control.

### Immunofluorescence

A549 cells seeded on coverslips were transfected with siRNA control or targeting JIP4 mRNA as described above. 48 hpt, cells were infected with 20 MOI of PR8. For synchronization of infection, cells were maintained at 4°C for 1 h followed by medium exchange and incubation at 37°C. Three hpi, cells were fixed with methanol for 10 min and maintained at 4°C in PBS. Blocking was performed with 3% BSA/PBS solution for 90 min on the shaker. 30 μl of a primary antibody mix consisting of mouse anti-NP (Bio-Rad MCA400) and rabbit anti-JIP4 (Cell Signaling Technology) diluted in 3% BSA were added and incubated for 1 h. Following three consecutive PBS washes, secondary antibodies (goat anti-rabbit IgG Alexa Fluor 488 from Invitrogen and goat anti-mouse Alexa Fluor 647 from Invitrogen) were added and incubated for 30 min protected from light. After three additional PBS washes, nuclei were stained with DAPI (Thermofisher) for 15 min protected from light. Coverslips were then washed with PBS and mounted for microscopy analyses. Pictures were taken with laser scanning microscope (LSM 780, ZeissOberkochen, Germany) equipped with Plan-Apochromat 63x (NA 1.4) oil immersion objective (Zeiss). All images were analyzed with ImageJ program.

## Results

### JIP4 presents anti-viral activity against IAV infection

Influenza virus replication leads to the regulation of several host pathways for both, viral benefits and host protective anti-viral responses. Different host proteins have been shown to act in tightly organized networks to assist in the activation or inactivation of specific pathways. JIP4 has not only been shown to bind to but also to interfere with MAPK JNK and p38 activation, two proteins highly reported to present relevant functions in IAV infection (7). In order to investigate a possible role of JIP4 in IAV infection, we transfected A549 cells with siRNAs specifically targeting JIP4 and analyzed the replication efficiency of IAV H1N1 PR8. Viral titers were increased in JIP4-silenced cells and this effect was maintained over multiple replication cycles resulting in significant differences 48 hpi **(Figure 1A)**. In agreement, overexpression of JIP4 resulted in a strong reduction in virus replication **(Figure 1B)**. Cells of the respective experiments were analyzed by Western blot to confirm the silencing or overexpression of JIP4. Similar to PR8, also WSN and the clinically relevant H1N1pdm09 strain presented higher replication rates in JIP4-silenced cells compared to control **(Figure 1C)**. Overall, these results suggest an anti-viral function of JIP4 in IAV infections.

**Fig 1.**
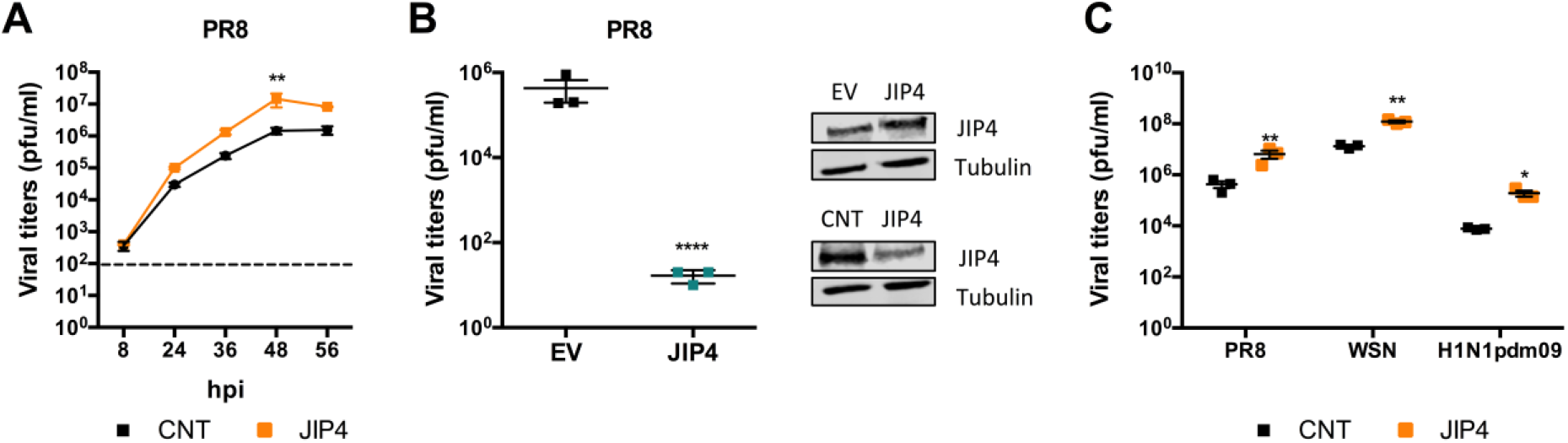
JIP4 presents an anti-viral activity against IAV infection. A549 cells were transfected with siRNAs targeting JIP4 or control siRNAs two days prior to infection. Cells were then infected with PR8 IAV (MOI 0.1). Supernatants were collected 8, 24, 36, 48 and 56 hpi and analyzed for viral titers by standard plaque assays. Cells were collected for evaluation of knock-down efficiency via Western blot, tubulin detection served as loading control. Results were statistically analyzed by two-way ANOVA test **(A)**. A549 cells were transfected with either, empty vector (EV) or plasmids containing JIP4 sequence, and infected with PR8 (MOI 0.1). Supernatants were collected 48 hpi for analysis of viral titers by standard plaque assay. Cells were collected for evaluation of overexpression efficiency by Western blot, tubulin detection served as loading control. Results were statistically analyzed by t-test **(B)**. A549 cells were transfected with siRNAs targeting JIP4 or control siRNAs and were infected with different IAV strains two days post transfection. PR8, WSN, and H1N1pdm09 were administered with 0.1 MOI. Supernatants were collected 48 hpi for determination of viral replication abilities by standard plaque assays. Results were analyzed by independent t-test per strain **(C)**. *p<0.01; **p<0.001; ****p<0.00001 compared to the respective control. All Western Blot images are representative of three independent experiments. Graphs are a compilation of three different experiments.

### JIP4 interferes with IAV replication prior to viral protein expression

We next examined at which step of the IAV life cycle JIP4 would restrict virus replication. Therefore, JIP4-silenced A549 cells were infected with PR8 MOI 5 and expression of viral proteins was analyzed by Western blotting **(Figure 2A)**. Interestingly, expression of viral proteins NS1 and PB1, which were chosen as representatives of early and late viral proteins respectively, was increased in the absence of JIP4 compared to control. To further confirm this phenotype, we evaluated viral protein amounts after overexpression of JIP4. Accordingly, viral protein expression was highly decreased in JIP4 overexpressing cells when compared to empty vector-transfected cells **(Figure 2B)**. Therefore, we conclude that JIP4 possesses an anti-viral activity already in early stages of viral replication, altering the production of viral proteins.

**Fig 2.**
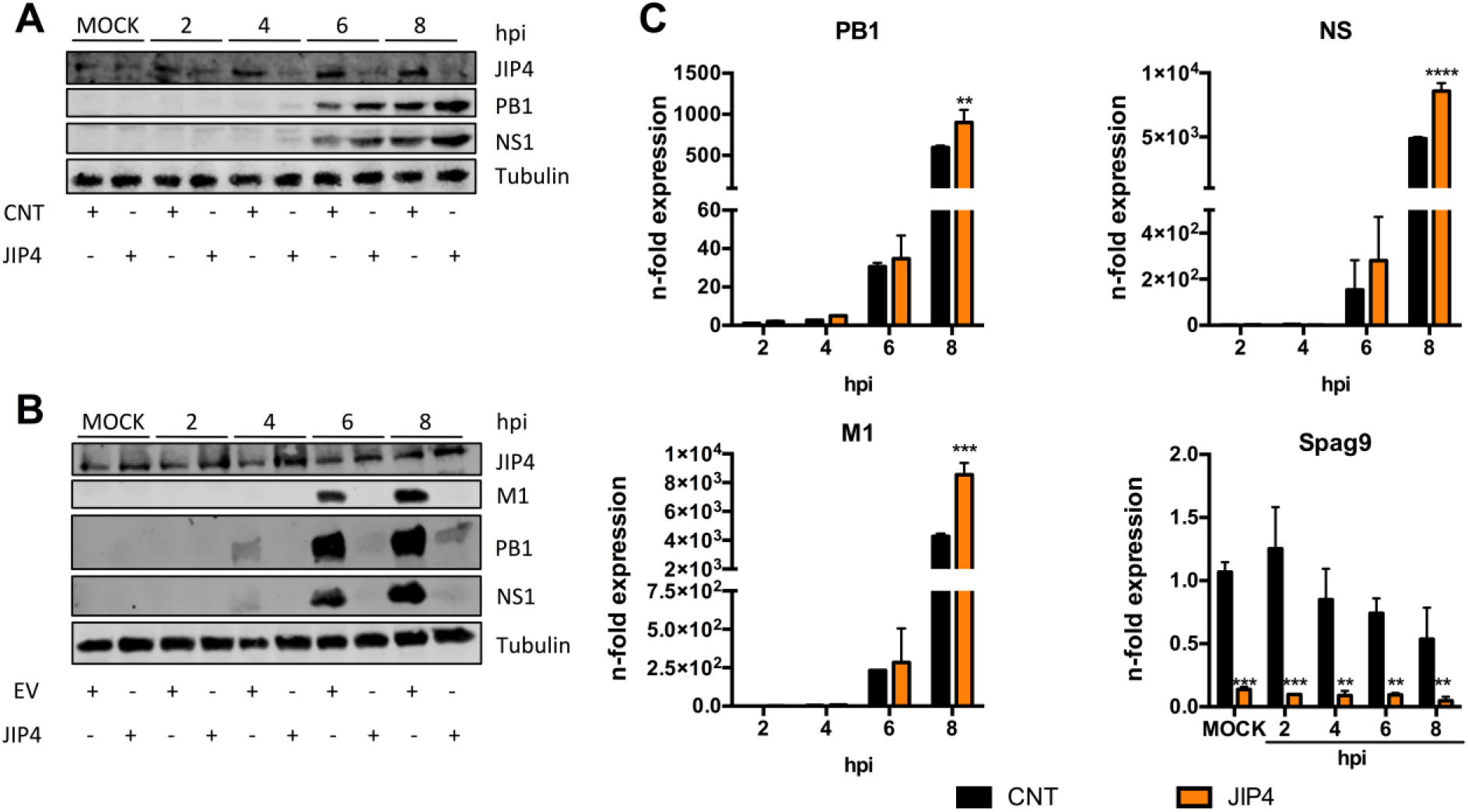
JIP4 interferes with IAV replication prior to viral protein expression. A549 cells were transfected with either, control siRNAs or siRNAs targeting JIP4, and infected with PR8 (MOI 5). Cells were lysed every 2 h. Lysates were submitted to Western blot assay and incubated with antibodies against viral proteins PB1 and NS1, as well as against JIP4 for silencing confirmation. Detection of tubulin served as loading control **(A)**. A549 cells were transfected with either, empty vector (EV) or plasmids encoding JIP4, and infected with PR8 (MOI 5). Cells were lysed every 2 h. Lysates were submitted to Western Blot assay and incubated with antibodies against viral proteins PB1, NS1 and M1, as well as against JIP4 for overexpression confirmation. Detection of tubulin was used as loading control **(B)**. For analyses of viral mRNA expression, A549 cells were transfected with either siRNAs control or targeting JIP4 and infected with PR8 (MOI 5). Samples were collected every 2 h and progressed for qRT-PCR analyses. Viral mRNAs were normalized over 2 h control group. *SPAG9* gene was normalized to MOCK control. Results were statistically analyzed by two-way ANOVA test **(C)**. **p<0.001; ***p<0.0001 and ****p<0.00001 compared to respective controls. All Western Blot images are representative of three independent experiments. Graphs are a compilation of three different experiments.

We next analyzed if the differences observed in viral protein expression are a consequence of alterations in translation or transcription steps by performing quantitative real-time PCR analysis of viral mRNAs PB1, NS and M1 **(Figure 2C)**. Interestingly, all viral mRNAs analyzed presented increased expression levels in JIP4-silenced cells when compared to control cells. Silencing efficiency was confirmed by the detection of low copy numbers of *SPAG9* mRNA.

Overall, these results demonstrate that the anti-viral function of JIP4 is effective in early stages of viral replication, resulting in altered viral mRNA levels and consequently viral protein expression, finally leading to decreased virus titers.

### The anti-viral function of JIP4 does not correlate with MAPKs JNK- or p38-induced anti-viral responses

In IAV infection, host cells produce interferons (IFNs) to induce an anti-viral state, which is mainly orchestrated by interferon beta (IFNß) (15). The presence of IFNs launches the expression of IFN-stimulated genes (ISGs) in neighboring cells which are subsequently protected from potential infection. Induction of IFNß expression is a result of the activation of different signaling cascades within the cell (16). Especially, our group has shown that activation of MAPKs p38 and JNK is required for type I IFN production in IAV infection (5, 6). Since JIP4 has been shown to bind to both, JNK and p38 MAPKs (7), we were prompted to evaluate if the anti-viral activity could be linked to a modulation of IFNß production. For that, A549 cells transfected with control or JIP4-targeting siRNAs, were infected with PR8 and analyzed for IFNβ and ISG expression by qRT*-*PCR. In control cells, a steadily raising induction of IFNβ on a low level correlated well with a continuous increase in expression of downstream ISGs MxA and IP10 upon infection **(Figure 3A)**. Interestingly, absence of JIP4 did not significantly alter the expression of those genes.

**Fig 3.**
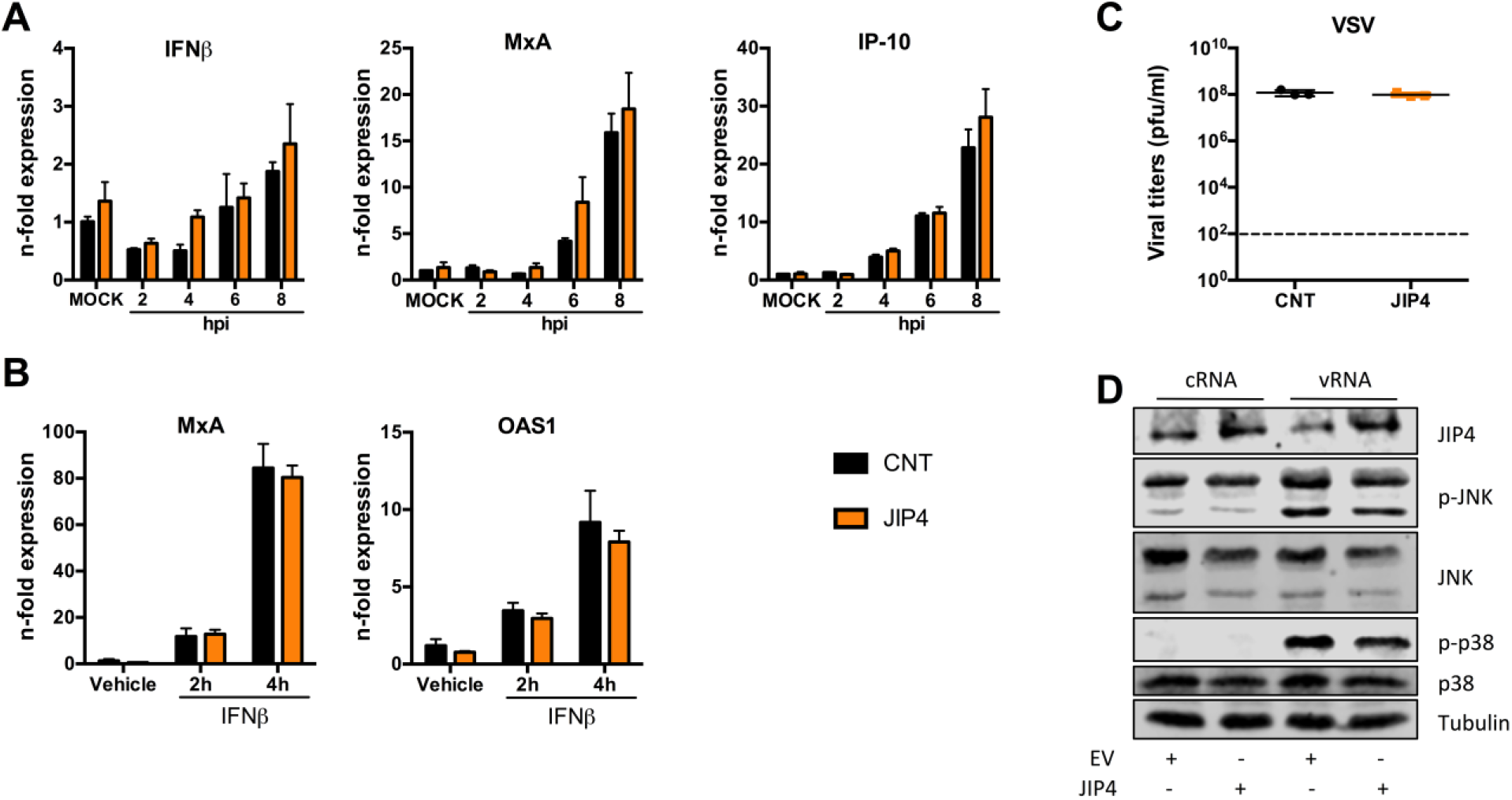
The anti-viral role of JIP4 does not correlate with changes in IFN responses or JNK/p38 activation. A549 cells were transfected with either, control siRNAs or siRNAs targeting JIP4, and infected with PR8 (MOI 5). Cells were collected every two hours for qRT-PCR analysis focusing on IFN and ISG mRNA expression. Results were statistically analyzed by two-way ANOVA test **(A)**. A549 cells were transfected with either, siRNAs control or targeting JIP4, and stimulated with IFNß for 2 or 4 h. ISG mRNA levels were analyzed by qRT-PCR. Results were statistically analyzed by two-way ANOVA test **(B)**. Results are depicted as n-fold expression over MOCK/vehicle after normalization to GAPDH expression **(A, B)**. A549 cells were transfected with either, siRNAs control or targeting JIP4, and infected with VSV (MOI 0.01). Supernatants were collected 24 hpi and virus replication was analyzed by standard plaque assays. Results are expressed in PFU/ml and statistically analyzed by t-test **(C)**. For analysis of JNK and p38 activity, A549 cells were transfected with either, EV or JIP4-expressing plasmids, and incubated for 24 h. Cells were then further transfected with c/vRNAs to stimulate activation of JNK and p38 kinases. After 3 h of stimulation, cells were lysed for Western blot analysis and activation of MAPKs was determined by using pJNK and p-p38 antibodies. Detection of p38, JNK and tubulin served as loading controls, successful overexpression was confirmed by JIP4 detection **(D)**. All Western Blot images are representative of three independent experiments. Graphs are a compilation of two independent experiments.

MAPK p38 has been shown to not only act in type I IFN production but also in IFN-mediated cell signaling (6). Therefore, we next evaluated if the absence of JIP4 would result in alterations in IFN signaling in a virus replication independent approach. For that, cells were transfected with siRNAs targeting JIP4 and stimulated with recombinant IFNß. qRT-PCR was used for quantification of ISG expression. MxA as well as OAS1 were induced upon stimulation, nevertheless, no differences in expression were observed between control and JIP4-silenced cells **(Figure 3B)**.

These results suggest that there is no involvement of IFN expression or response in the anti-viral activity of JIP4 in influenza A virus infection. In order to further confirm this hypothesis, we transfected cells with siRNAs targeting JIP4 and infected them with vesicular stomatitis virus (VSV) with an MOI of 0.1, which has been shown to be highly sensitive to IFN responses (17). As expected, the absence of JIP4 did not alter VSV replication **(Figure 3C),** clearly indicating that JIP4 does not function as a modulator of the type I IFN response.

Finally, since type I IFN production or signaling were not altered in absence of JIP4, we analyzed whether the protein would affect IAV-induced activation of kinases JNK and p38. For that, JIP4 overexpressing cells were stimulated with total RNA collected from either mock-infected (cRNA) or IAV-infected (vRNA) cells, and analyzed for the activation of MAPKs JNK and p38. In this assay, the vRNA pool isolated from infected cells serves as a pathogen associated molecular pattern (PAMP) to stimulate p38 and JNK activity. Our results clearly demonstrate that the overexpression of JIP4 does not significantly alter JNK or p38 activation after vRNA stimulation, even though both kinases showed a slight reduction in phosphorylation which can most likely be attributed to marginally decreased amounts of total protein after JIP4 overexpression **(Figure 3D)**.

Altogether, JIP4’s anti-viral role is neither correlated with an altered activation of JNK or p38 MAPKs nor with higher IFNβ responses.

### JIP4 interferes with viral polymerase activity

After receptor-mediated endocytosis of influenza virions and the release of the eight single-stranded viral RNA segments (vRNPs) into the cytoplasm (18), the trimeric viral RNA-dependent RNA polymerase, composed of PB1, PB2 and PA subunits, is responsible for the transcription and replication of the genomic segments (19). Since JIP4 altered the expression of viral mRNAs, we hypothesized that it might interfere with the machinery involved in viral RNA synthesis, hence, the viral polymerase. To test this hypothesis, we performed a mini-genome assay by transfecting cells with all subunits of the viral polymerase complex together with NP and an influenza-like luciferase reporter gene and either JIP4-encoding or empty vector plasmids. Viral polymerase reconstitution assays were performed in type-I IFN-deficient Vero cells in order to exclude any further interference of the type I IFN system. Indeed, JIP4-overexpressing cells showed a significant reduction in viral polymerase activity by approximately 75% when compared to control cells **(Figure 4A)**. We further analyzed whether viral genome replication is also modulated by JIP4 via evaluating the expression of three different RNA strands (vRNA, cRNA and mRNA) of gene segment 5 (NP) 4 hpi. In absence of JIP4, expression of all three RNA strands was significantly increased **(Figure 4B)**, which can most likely be attributed to an elevated expression of viral polymerase proteins already early in the replication cycle.

**Fig 4.**
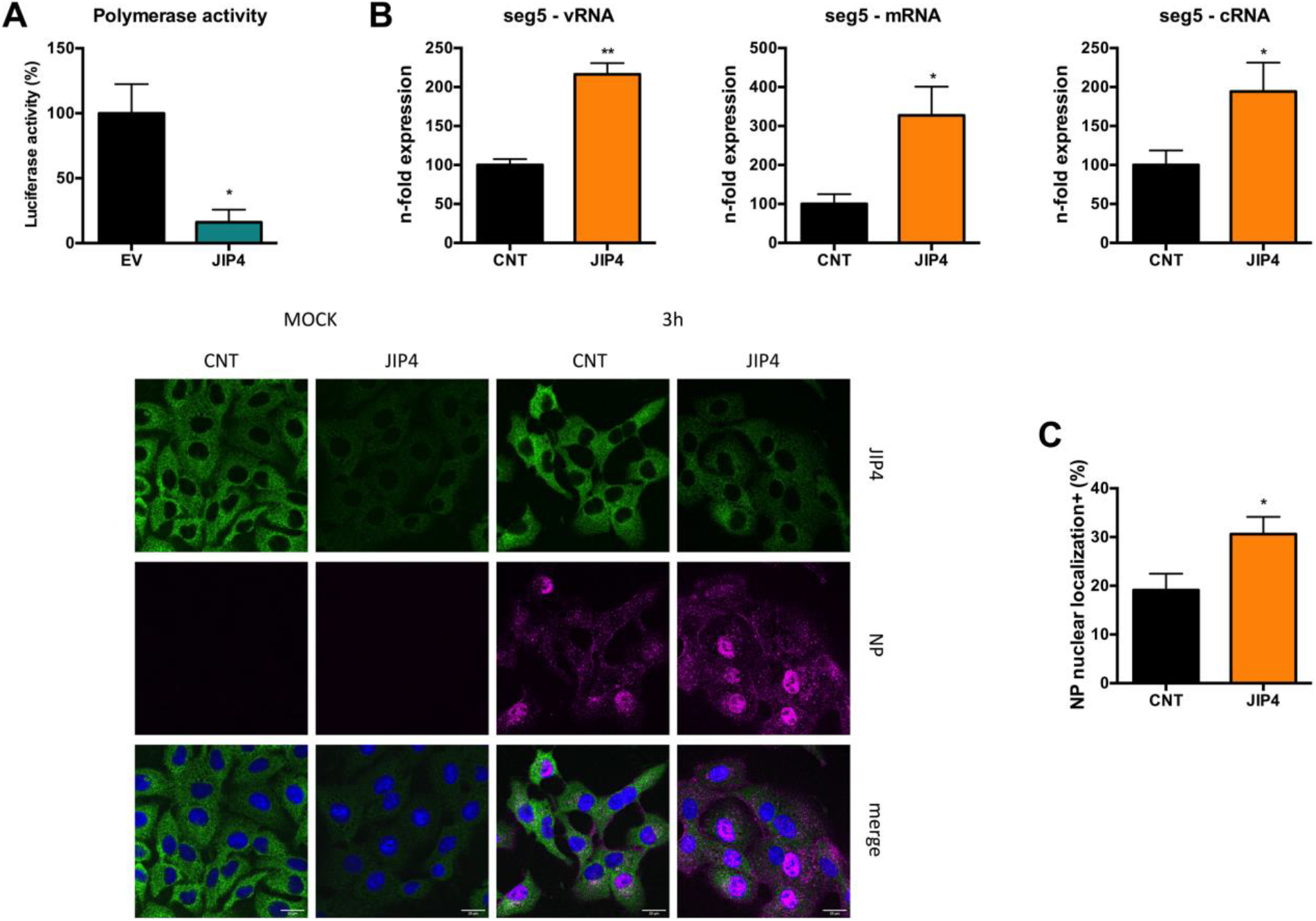
JIP4 overexpression decreased viral polymerase activity. Vero cells were transfected with either, EV or JIP4-expressing plasmids, together with all viral polymerase subunits and a reporter gene. 24 hpt cells were collected for luciferase assay analyses. Activity in EV-transfected cells was arbitrarily set to 100% **(A)**. A549 cells were transfected with either, siRNAs control or targeting JIP4, and infected with PR8 (MOI 5). Total RNA was collected 4 hpi for strand-specific qRT-PCR **(B)**. A549 cells were transfected with either, control siRNAs or siRNAs targeting JIP4. After 48 h of transfection, cells were infected with PR8 (MOI 20) and collected for immunofluorescence assay 3 hpi. Localization of JIP4 and NP was analyzed by using specific antibodies, nuclei were stained by DAPI. Percentage of cells with nuclear NP localization was expressed over the total number of cells in the field (**C)**. **p<0.001; *p<0.01 comparing to its respective control. Polymerase activity graph is a compilation of three independent experiments. Strand-specific PCR graphs are compilations of two independent experiments. Immunofluorescence image is representative of three independent experiments. All results above were statistically analyzed by t-test.

To understand whether JIP4 affects viral polymerase activity by a direct action on the polymerase complex in the nucleus, we analyzed the subcellular localization of JIP4 by immunofluorescence **(Figure 4C)**. As described previously (7), JIP4 is exclusively observed in the cytoplasm, even after infection. These results foreclose a direct interaction of JIP4 with the viral polymerase complex inside the nucleus. However, cells transfected with siRNAs targeting JIP4 presented a significantly higher percentage of NP-positive nuclei early in the replication cycle when compared to control cells.

Altogether, these results demonstrate that JIP4 possesses an anti-IAV activity that seems to be independent of a direct interference with viral polymerase complex formation in the nucleus but is most likely related to a hijacking activity with respect to incoming vRNPs as well as PB1-PA complexes, PB2 or NP in the cytoplasm.

### Phosphorylation of JIP4 at S730 is a prerequisite for its anti-viral function

Protein activities can be regulated by different mechanisms including posttranslational modifications such as phosphorylation or dephosphorylation. In order to analyze whether JIP4 function is modulated by dynamic phosphorylation in IAV infection, we performed a quantitative mass spectrometry analysis. A549 cells were infected with 5 MOI of IAV strains PR8, FPV or KAN-1, and JIP4 phosphorylation status as well as protein amounts were analyzed 0, 2, 4, 6 and 8 hpi. Respective phosphorylation intensities were normalized to JIP4 protein amounts and expressed as n-fold over non-infected cells. Here, we identified the serine at position 730 to be increasingly phosphorylated during viral life cycle progression. This phosphorylation was already detectable as early as 2 hpi with PR8 and KAN-1 strains and further elevated after infection with all tested strains 6 hpi **(Figure 5A)**. Next, we wanted to investigate if S730 phosphorylation is responsible for the anti-viral activity of JIP4 in IAV infection. Therefore, we used plasmids containing mutations in the specific phosphosite changing the serine to a non-phosphorylatable alanine (A) or glutamic acid (E), mimicking a constitutively phosphorylated state, and analyzed IAV polymerase activity. As observed previously, overexpression of wild type (WT) JIP4 resulted in a significant reduction in viral polymerase activity. Interestingly, presence of non-phosphorylatable JIP4 S730A rescued the activity observed in empty vector-transfected control cells. In contrast, JIP4 S730E reduced viral polymerase activity to a similar level as observed for WT JIP4 **(Figure 5B)**. These results indicate a possible relationship between JIP4 S730 phosphorylation and the anti-viral role of the protein in IAV infection.

**Fig 5.**
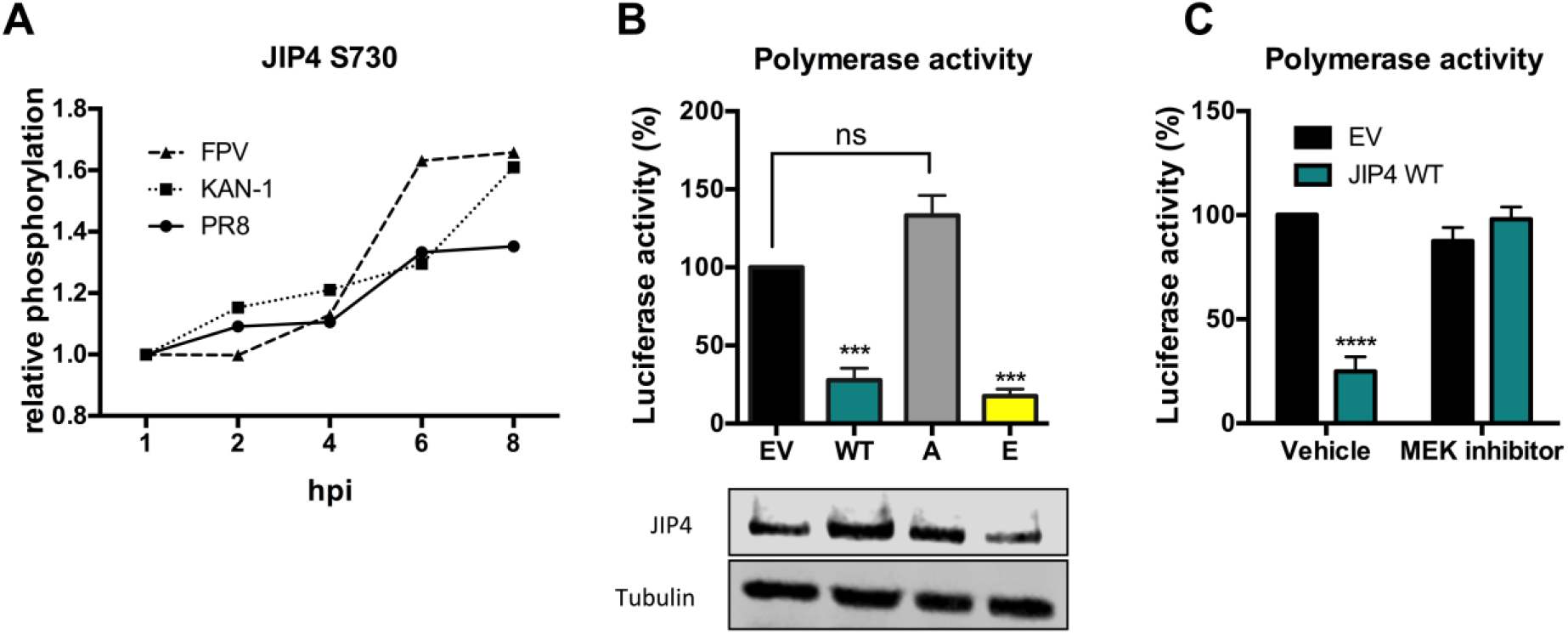
Phosphorylation of JIP4 S730 results in a decreased viral polymerase activity. A549 cells were labeled by using SILAC and infected with low pathogenic IAV strain PR8 (H1N1) or one of two highly pathogenic strains (H5N1 – KAN-1 and H7N7 - FPV). Samples were collected 2, 4, 6 and 8 hpi and analyzed for JIP4 phosphorylation. Relative phosphorylation of JIP4 at S730 at different time points compared to non-infected cells is depicted after normalization to respective JIP4 protein amounts **(A)**. Vero cells were transfected with plasmids expressing either, EV, WT JIP4, or JIP4 phospho-mutants S730A or S730E, together with all viral polymerase subunits and a reporter gene. 24 hpt, cells were collected for luciferase assay analysis. Activity in EV-transfected cells was arbitrarily set to 100% and all groups were statistically compared with EV-transfected cells by one-way ANOVA test. Overexpression was confirmed by Western blot **(B)**. Vero cells were transfected with either, EV or plasmids expressing WT JIP4, together with all viral polymerase subunits and a reporter gene. After 4 h of transfection, medium was replaced by fresh medium supplemented with vehicle or 10 μM MEK inhibitor U0126. Luciferase assay was performed 24 hpt. Activity in EV-transfected cells group was arbitrarily set to 100%. All groups were compared by two-way ANOVA test. **(C)**. ***p<0.0001 and ****p<0.00001 compared to control group. Mass spectrometry phosphorylation kinetic data is a result of one single experiment. Both polymerase activity graphs are a compilation of three independent experiments.

Next, we decided to analyze the cellular signal transduction pathway that might be responsible for JIP4 phosphorylation at S730. It has already been reported that different kinases present important roles during IAV infection and MAP kinases, especially the Raf/MEK/ERK pathway, have been previously reported to be involved in JIP4 S730 phosphorylation regulation (20, 21). Therefore, we analyzed whether the Raf/MEK/ERK pathway could be responsible for JIP4 phosphorylation and, consequently, for its anti-viral function. Viral polymerase activity was evaluated in presence of WT JIP4 with or without MEK inhibitor U0126. Interestingly, as observed previously, WT JIP4 strongly decreased viral polymerase activity **(Figure 5C)**, whereas addition of the MEK inhibitor resulted in no difference between empty vector- and WT JIP4-transfected cells. Therefore, we can conclude that the Raf/MEK/ERK pathway is involved in the phosphorylation of JIP4 orchestrating its anti-viral activity.

## Discussion

IAV is a widely spread respiratory pathogen that infects 10% of the global population annually. Anti-viral treatments are urgently needed. Targeting host pathways is a promising approach to avoid development of resistance. We have identified a novel anti-viral function of the cellular JIP4 protein in IAV infection that is independent of type I IFN production or p38/JNK MAPK activation. JIP4 influences the IAV life cycle in early stages of viral replication, resulting in decreased viral polymerase activity. Furthermore, we identified the phosphorylation of JIP4 at S730, which is mediated by the Raf/MEK/ERK pathway, to be decisive for its anti-viral function.

We demonstrate an interesting pattern of viral JIP4 dependency, in which the lack of JIP4 resulted in higher viral replication of IAV. Our results also point to an early effect in viral replication, since we could already observe differences in viral mRNA and protein expression as early as 4 hpi.

As already suggested by the name of the protein family, one of the well-known interaction partners of the JIP family is JNK, since all members possess a JNK binding site (22). JNK activation has been observed after infection with several different viruses, and, particularly in IAV infection, activation of JNK was shown to exhibit both, virus-supportive and anti-viral functions (23–25). Besides this common function of JIP family members, JIP4 has specifically been reported to also bind to p38 MAPK (8). The link between the p38 MAPK pathway and inflammation has been well established. Our group already reported how p38 inhibition leads to protection against IAV *in vivo* due to its suppression of cytokine amplification. In addition, our group also showed the influence of p38 inhibition on the production of main antiviral responses orchestrated by type I IFN, and especially IFNß (6). In accordance with the literature, JIP4 overexpression resulted in slight alterations of both, phosphorylated and total amounts of JNK and p38. Nevertheless, the absence of JIP4 did not result in alterations in the production of IFN or ISGs. These results are in agreement with the literature which reported key differences of JIP4 when compared to other proteins of the JNK-interacting protein group. The lack of ability to bind to MAP kinase kinase 7 (MKK7) and Mixed lineage kinase 3 (MLK3), two upstream proteins of the JNK signaling pathway, suggests that JIP4 might not act as a scaffold protein as already characterized for the other members of the group (7). In this sense, we could exclude a modulation of p38 and JNK activation as an explanation for the observed anti-viral activity of JIP4.

JIP4 absence resulted in lower viral mRNA levels, demonstrating interference with the viral life cycle prior or during transcriptional steps. IAV polymerase activity showed a significant decrease upon JIP4 overexpression without an indication of a direct interference with the viral polymerase complex in the nucleus. Since JIP4 is a recently discovered protein, little is known about its different roles in cell biology and especially during viral infections. Hence, it is still mechanistically unclear how JIP4 alters viral polymerase activity. JIP4 has previously been shown to interact with ARF6 (ADP-ribosylation factor 6) at the cell membrane, which was correlated with the control of trafficking of recycling endosomes in and out of the intercellular bridge, together with the function in abscission during cell division (9). Nevertheless, our minigenome assay is performed with direct transfection of the polymerase complex subunits, thus, we can already exclude that JIP4 would alter viral entry and/or uncoating, both early steps in viral replication that depend on host cell membranes. The exact mechanism of how JIP4 interferes with IAV replication remains elusive, but we hypothesize that JIP4 interferes with the import of incoming vRNPs or distinct newly synthesized polymerase subunits into the nucleus.

The anti-viral activity of JIP4 is correlated with S730 phosphorylation mediated by MEK, ERK or a further downstream acting kinase. The site S730 has already been identified to be phosphorylated in different species including humans, mice and rats. Nevertheless, no specific role has been assigned to it, yet. The importance of the virus-supportive role of the Raf/MEK/ERK signaling cascade in IAV infection is well documented (26–28). ERK signaling has been shown to be activated in different stages of the IAV life cycle, including the first hours during viral entry and uncoating as well as a second activation during vRNP nuclear export (29, 30). Interestingly, previously described ERK activation dynamics nicely correlate with the observed JIP4 S730 phosphorylation kinetics, since it seems to be already in a phosphorylated state 2 hpi and further increased after 6 h of IAV infection. Our group has exceedingly shown that MEK inhibition results in an overall decreased viral replication (26, 28), highlighting again how efficient viruses are in the misuse of cellular pathways that also provide anti-viral functions as observed previously for MAPK JNK or the NFκB pathway (4, 31, 32). Further unraveling of the detailed mechanism of the anti-viral activity of JIP4 might provide alternative ways to develop new anti-viral approaches.

In summary, we have identified a novel anti-viral function of the JIP4 protein in IAV infection. We showed that the virus-restricting function of JIP4 is not due to a modulation of JNK or p38 MAPK signaling cascades or alterations in IFN production or signaling. We have demonstrated that JIP4 alters viral polymerase activity without direct interaction with the vRNP complex in the nucleus. Furthermore, we have identified dynamic phosphorylation of JIP4 at S730 mediated by the Raf/MEK/ERK kinase cascade, to be responsible for the observed anti-viral function. We conclude that JIP4 might be a promising target for IAV antiviral therapy.

## Acknowledgements

We thank Saskia Hinse and Felicia Ye Wen Hwa for excellent technical assistance. This work was supported by the German Research Foundation (DFG, grants Lu477/25-1, BO5122/1-1, RTG 2220 A1 and SFB1009 TPB02), the Interdisciplinary Centre for Clinical Research (IZKF, Lud2/017/13) and CAPES with PDSE scholarship. The funders had no role in study design, data collection and interpretation, or the decision to submit the work for publication.

